# Advancement of spring arrival and delayed autumn departure in Ruby-throated Hummingbird (Archilochus colubris) migration phenology over more than a century

**DOI:** 10.64898/2026.09.03.749133

**Authors:** Sydney M. Pierce, Terra M. Swartz

## Abstract

Climate change is thought to have altered the timing of avian migration, with species-specific responses varying across spatial and temporal scales. This is notable as ecological mismatches may occur when birds arrive or depart out of sync with the peak flowering period of their associated food plant resources. This study compares first arrival and last departure dates of *Archilochus colubris* (Ruby-throated Hummingbird) using citizen science data from the North American Bird Phenology Program for historical records (1880-1969) and recent records from eBird (2006-2024) and Journey North (2001-2024). Results show significantly earlier spring arrivals (14.2-27.9 days) and later fall departures (17.3-33.7 days) in the recent period compared to historical periods, while migration pace remained relatively consistent across datasets. Although temperatures are increasing in both the nonbreeding and breeding ranges of *A. colubris*, winter and fall temperatures were not strong independent predictors of migration timing. This study is the first to document shifts in fall departure dates for the species and highlights the need for continued research to uncover the mechanisms driving these phenological changes and assess their broader ecological implications.

**LAY SUMMARY:**

- *Archilochus colubris* (Ruby-throated Hummingbird) now arrive earlier in the spring and depart later in autumn across the United States compared to historical patterns.
- These changes in migration timing have occurred over more than a century.
- Despite changes in arrival and departure dates, the overall speed of migration has stayed moderately the same.
- Changes in seasonal temperature alone do not fully explain these long-term migration shifts.
- Altered timing may affect how hummingbirds interact with flowering plants and could have broader impacts on ecosystems.

## INTRODUCTION

The year 2024 was the warmest on record, with global temperatures rising 1.6 °C above pre-industrial levels (Bevacqua et al. 2025). Bird migratory schedules are often used to examine how anthropogenic climate change is affecting the natural world, as monitoring programs have been in place for more than a century (Zelt and Droege 2010; Wilson 2013; Courter 2017). Recent studies have reported that a range of migratory hummingbird species have earlier arrival dates than previous periods (McKinney et al. 2012; Probst et al. 2017; Courter 2017). This phenomenon is thought to be largely driven by changes to the global climate, as weather variables such as temperature and precipitation are predictors of avian migration timing (Kelly et al. 2016; Hinchcliffe and Tkaczynski 2025). However, non-climate mechanisms may also be affecting avian migration phenology, such as earlier flowering onset, as hummingbirds have an innate urgency to arrive at breeding grounds when resources are plentiful (Correa-Lima et al. 2019; Geissler et al. 2023).

Hummingbirds have a specialized mutualistic relationship with their associated food plant species, characterized by complementary morphological traits. Trait matching is generally prevalent between plant corollas and hummingbird mouthparts (Maglianesi et al. 2014; Leimberger et al. 2022; Maglianesi et al. 2024). This increases the efficiency of nectar extraction while simultaneously ensuring that plants receive minimal cross-pollination from non-specialist species (Klumpers et al. 2019; Rico-Guevara et al. 2021). Specialization is believed to drive speciation, facilitate resource partitioning, and enhance ecosystem functioning (Phillips et al. 2020; Leimberger et al. 2022). Thus, temporal mismatches between the arrival of hummingbirds and the peak flowering period of their floral species could have significant ecological consequences. These may include co-extinction, reduced reproductive success, the decoupling of plant-hummingbird interactions, and the inability of other pollinators to compensate for the absence of hummingbirds (Hadley et al. 2014; Remolina-Figueroa et al. 2022; Ponti and Sannolo 2023; Trunschke et al. 2024). Potential repercussions emphasize the importance of understanding migratory changes so to implement conservation efforts.

Bird migratory data is rapidly becoming available through the development of the citizen-science projects eBird (eBird.org) and Journey North (journeynorth.org). Users of these sites can submit bird observations via the Internet to a centralized data repository. These efforts have allowed a better understanding of avian migratory patterns, abundance, and distribution (Kelly et al. 2016; Walker and Taylor 2017; Horns et al. 2018; Johnston et al. 2021). While this makes eBird and Journey North excellent sources for tracking recent avian migration patterns, they alone cannot be used to assess the impact of global warming due to limited data availability prior to 2001 (Courter 2013; Sullivan et al. 2014).

The North American Bird Phenology Program (NABPP), a lesser-known counterpart to the contemporary citizen science projects, is the longest and most detailed legacy dataset on North American bird migration (Zelt et al. 2012). Started in 1881, the program recorded more than 870 North American bird species first arrival dates, departure dates, and populations through the efforts of naturalist volunteers (Zelt and Droege 2010; Zelt et al. 2012). Although the program ceased in 1970 and was largely forgotten, it was revitalized in 2008 by the U.S. Geological Survey in response to growing concerns about how climate change affects bird phenology (Zelt and Droege 2010; Zelt et al. 2012). Now, over 6 million migration card observations have been transcribed and digitized, providing a unique historical baseline for bird migration research (Zelt and Droege 2010). When combined with eBird and Journey North for recent observation records, the NABPP offers immense potential to evaluate the impacts of climate change on bird migration across broad spatial and temporal scales.

*Archilochus colubris* (Ruby-throated Hummingbird) is an abundant migratory species that breeds in the eastern United States (Robinson et al. 1996; Osborne 1998). In the spring, these hummingbirds travel northward from their wintering grounds in Central America to their breeding territories, which span the East Coast, the central Great Plains, and extend into southern Canada and the Gulf Coast (Robinson et al. 1996; Osborne 1998). Males typically migrate ahead of females, reaching southern states by mid to late March and northern regions between late April and early May (Goodrich 1988; Robinson et al. 1996; Osborne 1998). Between August and October, these Neotropical migrants vacate their breeding territories and depart south to Central America (Goodrich 1988). Both spring and fall migration for *A. colubris* is synchronous with the flowering periods of their primary nectar sources, which include floral species found in mixed woodlands and gardens, such as *Impatiens capensis* (jewelweed) and *Lobelia cardinalis* (cardinal flower) (Goodrich 1988; Osborne 1998). Thus, their migratory patterns are crucial in sustaining their high metabolic demand and facilitating pollination.

Given their distinctive plumage and their presence in both suburban and urban areas, there is an extensive record of *A. colubris* observations available to the public. Recent studies have utilized this data and found that the changing climate has influenced earlier arrival times to breeding grounds when compared to previous periods (Courter et al. 2013). However, to our knowledge, there are no studies examining the changing phenology of hummingbird departure dates during fall migration. Therefore, using the available geographic databases of observations, we assess the changes in both arrival dates and departure dates in *A. colubris* between historical (1880-1969) and recent periods (2001-2024) in the United States in relation to temperature.

## METHODS

### Arrival and Departure Data

All data analysis and visualization were performed using R version 4.5.3 (R Core Team 2025). Historical migration records of *A. colubris* (1880-1969) were obtained from the NABPP (Droege et al. 2023), while recent migration records were sourced from eBird (2006-2024) and Journey North (2001-2024) through their respective online platforms. Although both eBird and Journey North contain records prior to these dates, data from earlier years were sparse and geographically limited. Therefore, 2001 and 2006 were selected as starting points due to a marked increase in observation frequency and data quality from these years onward (Courter et al. 2013; Sullivan et al. 2014; Courter 2017). Journey North data were filtered to exclude entries with missing values or zero counts, and eBird data were restricted to include only complete checklists that reported all species observed during an observation period to enhance data accuracy. To facilitate spatial analysis, a one-degree resolution spatial grid was constructed that encompassed North America (Figure 1). All observations were standardized to an ordinal date format (e.g., April 10 = day 100), with adjustments made for leap years, and were spatially joined to the grid based on observation latitude and longitude. Grid cells with fewer than 100 eBird observations total were excluded to ensure adequate sample sizes.

**Figure 1.**
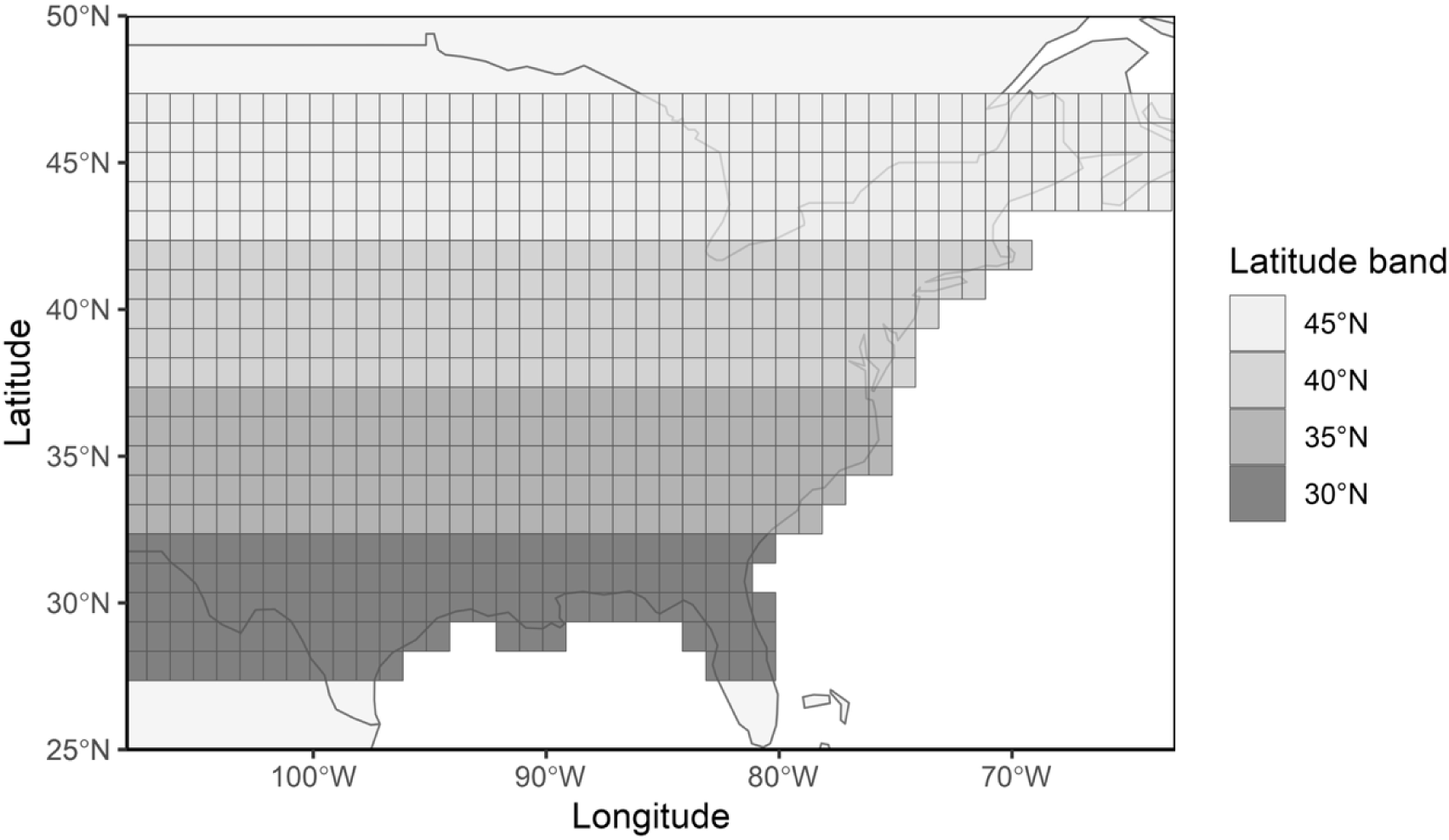
One-degree spatial grid used to standardize migration analyses across North America. Each grid cell represents a 1° latitude × 1° longitude spatial unit to which individual observations of *A. colubris* were assigned throughout the species’ migratory range. Shading indicates the 4 5° latitudinal bands used in the analyses. Alt text: Map of North America overlaid with a 1° latitude and longitude grid spanning the breeding range of *A. colubris*. Grid cells are grouped into 4 latitudinal bands (30-32°N, 32-37°N, 37-42°N, and 42-47°N), referenced as 30°, 35°, 40°, and 45°N. Axes display latitude (25-50°N) and longitude (108-63°W).

Arrival dates were defined as the earliest observation recorded in each year for each grid cell for all datasets. Departure dates were defined as the last recorded observation per year within each grid cell, using only the NABPP and eBird data, as Journey North does not provide departure information. Observations north of 45°N and south of 30°N bands were excluded from analysis due to low data density. Additionally, to eliminate the influence of potential sedentary or vagrant birds, only observations within biologically relevant migration windows were used, with arrival data restricted to February 1-May 31 and departure data restricted to June 1-October 31.

To assess spatiotemporal trends in migration timing and detect differences across datasets, arrival and departure data were aggregated by latitude, decade, and data source. Data source is a critical variable because the NABPP data were collected from 1880 to 1969, representing historical trends in migration, prior to the effects of anthropogenic climate change. In contrast, eBird (2006-2024) and Journey North (2001-2024) data represent modern migration trends under the effects of climate warming, allowing for a comparison of migration timing before and after substantial climate shifts. The influence of latitude and the data source on arrival and departure dates was evaluated using a two-way analysis of variance (ANOVA) with latitude band and data source as fixed factors. The interaction between these variables was also tested to determine whether the effect of latitude on migration timing varied across data sources. Pairwise comparisons were performed using Tukey’s Honest Significant Difference (HSD) test to identify significant differences in migration dates between latitudes and data sources. The mean number of days between successive 5° latitude bands was also calculated from the mean first arrival and last departure dates at each latitude, providing a proxy for migration pace.

### Climate Data

Temperature data were obtained from the Climatic Research Unit Time-Series dataset (CRU TS* 4.09) accessed via the Centre for Environmental Data Analysis archive (University of East Anglia Climatic Research Unit et al. 2025). Developed by the University of East Anglia, CRU TS is a globally gridded dataset with a spatial resolution of 0.5° × 0.5° that provides monthly surface air temperature estimates from January 1901 through December 2024. The dataset is derived from observations collected at more than 4,000 meteorological stations worldwide, which are spatially interpolated through angular-distance weighting to generate continuous fields describing monthly temperature variability (Harris et al. 2020).

To evaluate the influence of winter climate on spring migration phenology, temperature data were extracted from grid cells corresponding to the nonbreeding range of *A. colubris* in Central America. January and February were selected to represent late-winter conditions that precede migration. For each year (1901-2024), mean monthly temperatures were calculated by averaging values across all grid cells within the defined region, and a winter temperature index was computed as the mean of January and February temperatures. These temperature data were subsequently joined with annual mean first arrival dates, with arrival records restricted to observations representing mean arrival timing within spatial grid cells in the 30°N latitude band. This approach was motivated by a previous study demonstrating that earlier spring arrival in *A. colubris* may be associated with elevated temperatures in wintering grounds, reflecting the potential influence of pre-migratory climatic conditions on migration initiation (Courter et al. 2013).

For statistical analysis of winter climate effects on arrival timing, a set of a priori linear regression models was constructed with first arrival day as the response variable. Candidate models represented hypotheses that arrival timing was explained by temporal change, February temperatures on nonbreeding grounds, mean winter temperatures on nonbreeding grounds, or combined temporal and climatic effects. Competing models were compared using second-order Akaike’s Information Criterion corrected for small sample sizes (AICc), and model support was assessed using ΔAICc and Akaike weights (*w_i_*). Collinearity among predictors was evaluated using variance inflation factors (VIF).

To examine the influence of fall climate conditions on departure timing, temperature data were extracted from grid cells corresponding to the breeding range of *A. colubris* across all 4 latitude bands included in this study (30°, 35°, 40°, and 45°N). Monthly temperatures were obtained for June through October to represent the late breeding and fall migration period (Goodrich 1988). For each latitude band and year, mean temperatures were calculated for each month by averaging grid cell values within each spatial extent. A seasonal fall temperature metric was then calculated as the mean temperature across the June-October period. These data were joined with annual mean last departure dates for each latitude band, allowing evaluation of whether *A. colubris* exhibits sensitivity to temperature anomalies through local environmental cues and thus experiences fall migration shifts.

For analysis of fall climatic effects on departure timing, a separate set of a priori linear regression models was constructed with last departure day as the response variable. Candidate models represented hypotheses that departure timing was explained by temporal change, mean fall temperatures on breeding grounds, latitude band, and their combined or interactive effects. Competing models were evaluated using AICc, and relative model support was assessed using ΔAICc and *w_i_*to identify the most parsimonious explanation for variation in departure timing.

## RESULTS

### Arrival Data

Comparisons using Tukey’s HSD test showed that mean first arrival dates differed significantly between historical and recent periods across latitudes (Supplementary Materials Table S1), with *A. colubris* arriving 14.2-27.9 days earlier (Table 1). Differences in arrival dates between the NABPP and eBird decreased with increasing latitude, while differences between the NABPP and Journey North were less pronounced at the 30°N and 45°N latitude bands (Table 1). Additionally, migratory rates increased at higher latitudes for both the NABPP and eBird (Table 2). The mean difference in arrival dates between eBird and Journey North was negligible (*P* = 0.99), indicating no significant difference in migration timing between these recent data sources. The two-way ANOVA revealed significant effects of both latitude (F(3, 15779) = 12548.9, *P*<0.001) and data source (F(2, 15779) = 4530.9, *P*<0.001) on first arrival dates, as well as a significant interaction between the 2 factors (F(6, 15779) = 113.5, *P*<0.001). These results are further supported when visualized by decade and data source, which shows a clear trend of earlier first arrival dates in the recent decades compared to historical decades, while migration rates remain similar across sources and decades (Figure 2).

**Figure 2.**
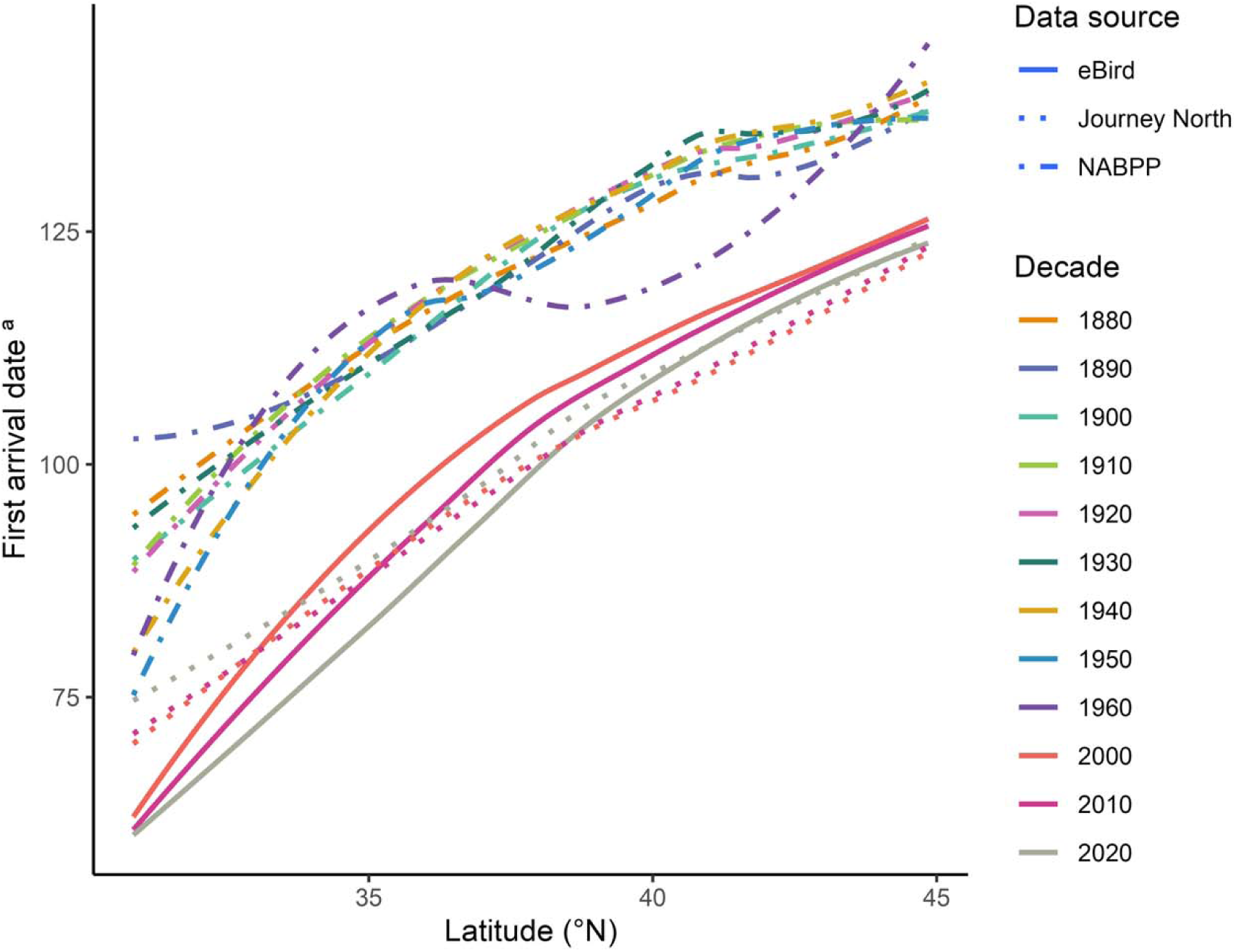
First arrival dates of *A. colubris* across latitudes, with trends depicted by decade and data source. ª Arrival dates expressed as day-of-year and corrected for leap years. Alt text: Line graph showing first arrival date of *A. colubris* across latitude bands 30°N to 45°N. Colored lines represent decades spanning the 1880s to 2020s, and line types distinguish eBird, Journey North, and NABPP datasets. Arrival dates have advanced in recent decades (2000s-2020s), but the pace of migration has remained relatively consistent.

**Table 1.**
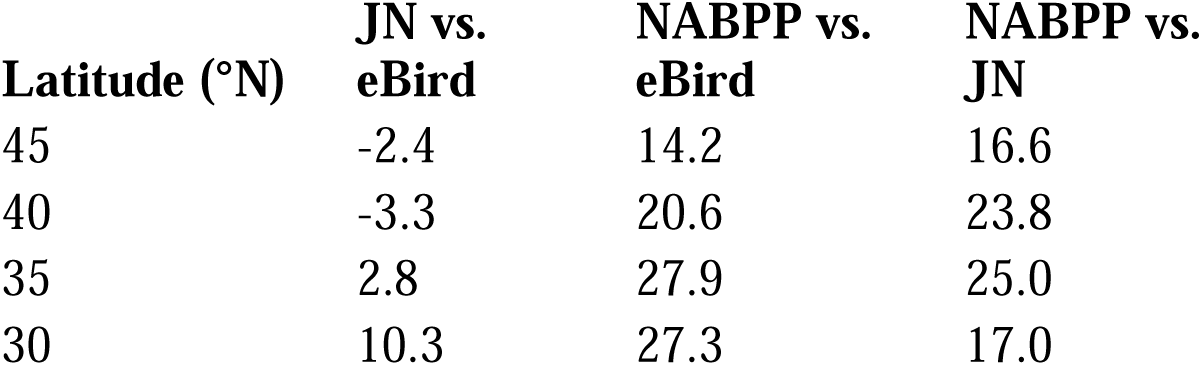
Differences in *A. colubris* mean first arrival day between recent data sources, eBird and Journey North (JN), and historical data source, the NABPP, across 4 latitude bands. Positive values indicate later arrival in the first-named source, while negative values represent later arrival in the second-named source.

**Table 2.** Mean spring migration pace of *A. colubris* between successive 5° latitude bands from recent data sources, eBird and Journey North, and historical data source, the NABPP. Values represent the interval in days between mean first arrivals at each band, providing a proxy for migration pace.

| Latitude (°N) | eBird | Journey North | NABPP |
| --- | --- | --- | --- |
| 30 → 35 | 21.4 | 14.0 | 22.0 |
| 35 → 40 | 25.7 | 19.6 | 18.4 |
| 40 → 45 | 11.7 | 12.6 | 5.4 |

### Departure Data

*A. colubris* departure dates also differed dramatically between historical and recent periods. Tukey’s HSD test indicated that mean departure dates were 17.3-33.7 days later in the recent period, with the smallest difference observed at the 30°N latitude band (Table 3). Additionally, fall migratory pacing differed notably between 35°N and 30°N, with birds requiring a mean of 19 days to reach the 30°N band in the NABPP dataset, compared to just 2 days in the eBird dataset. (Table 4). Differences in departure timing between northern latitude bands were not significant. The ANOVA again revealed significant effects of both latitude (F(3, 5237) = 261.9, *P*<0.001) and data source (F(1, 5237) = 2443.9, *P*<0.001) on departure dates, as well as a significant interaction between the 2 factors (F(3, 5237) = 6.4, *P*<0.001). While trends in last departure dates are less pronounced than those in arrival dates when visualized by decade and data source, the data still indicates a general pattern of later departures in recent decades (Figure 3).

**Figure 3.**
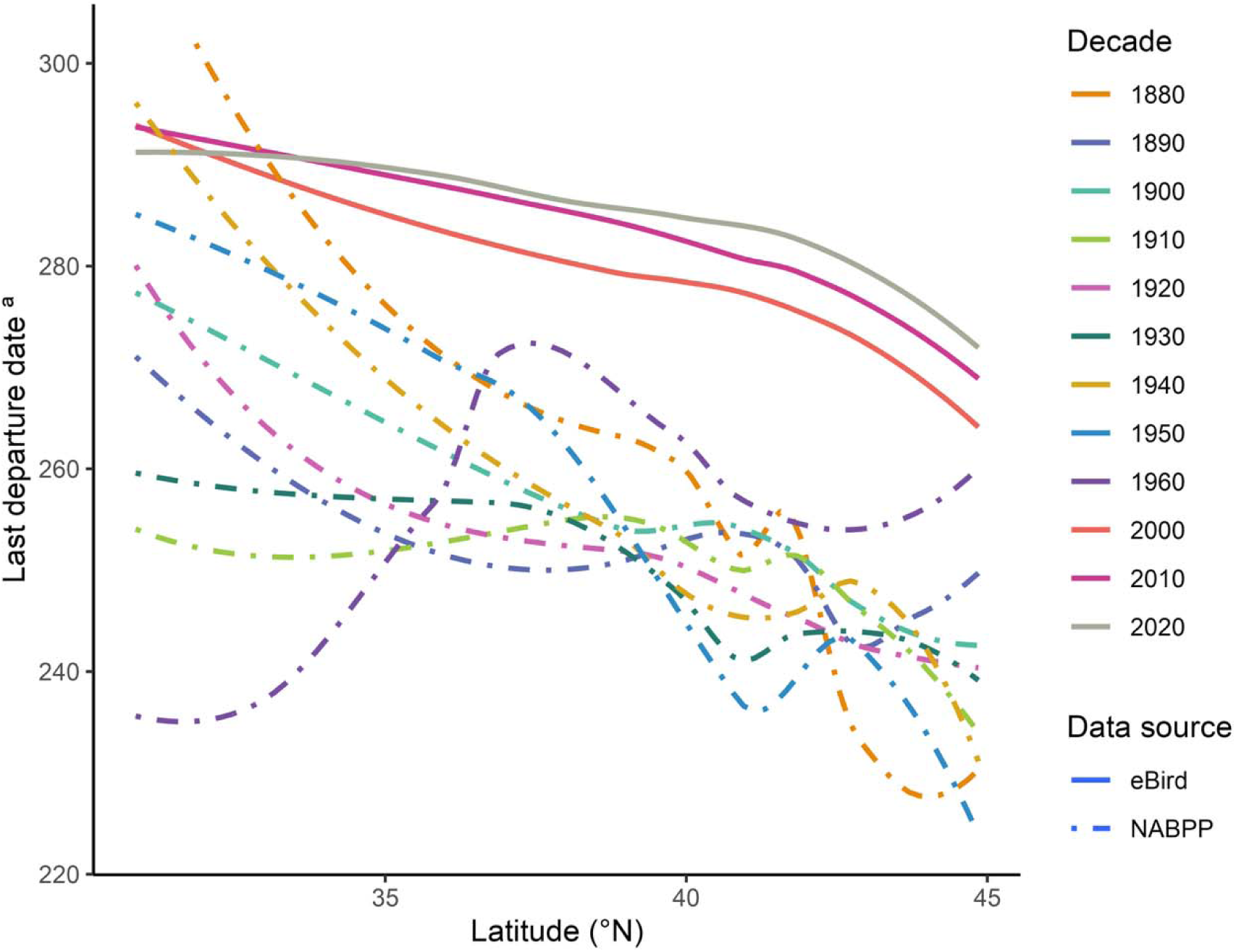
Last departure dates of *A. colubris* across latitudes, with trends depicted by decade and data source. ª Departure dates expressed as day-of-year and corrected for leap years. Alt text: Line graph showing last departure dates of *A. colubris* across latitude bands 30°N to 45°N. Colored lines represent decades spanning the 1880s to 2020s, and line types distinguish the eBird and NABPP datasets. Departure dates occur later in more recent decades (2000s-2020s), indicating a shift toward extended fall residence time across the breeding range.

**Table 3.** Differences in *A. colubris* mean last departure day between the recent data source, eBird, and historical data source, the NABPP, across 4 latitude bands. Negative values indicate later departure in the second-named source, eBird.

| Latitude (°N) | NABPP vs. eBird |
| --- | --- |
| 45 | -31.9 |
| 40 | -32.2 |
| 35 | -33.7 |
| 30 | -17.3 |

**Table 4.** Mean fall migration pace of *A. colubris* between successive 5° latitude bands from recent data source, eBird, and historical data source, the NABPP. Values represent the interval in days between mean last departures at each band.

| Latitude (°N) | eBird | NABPP |
| --- | --- | --- |
| 45 → 40 | 8.4 | 8.0 |
| 40 → 35 | 6.5 | 5.1 |
| 35 → 30 | 2.6 | 19.0 |

### Climate Data

Model selection indicated that arrival timing was best explained by year alone, supporting a trend toward earlier spring arrival over time (Table 5A). Interestingly, models including temperature variables as a sole predictor received little support. Models incorporating year with February or winter temperature were competitive but did not substantially improve model fit relative to the year-only model. Variance inflation factors were low (VIF<2.5), indicating that multicollinearity among predictors was minimal.

**Table 5.** Model selection results evaluating a priori hypotheses for predictors of migration phenology. Candidate models were ranked using second-order Akaike’s Information Criterion corrected for small sample size (AICc). For first arrival timing (**A**), the lowest AICc was 9985.14; for last departure timing (**B**), the lowest AICc was 44399.90. *k* = number of estimated parameters; Dev. = deviance (−2lnL); ΔAICc = difference from the best-supported model; *w_i_* = Akaike weight; Cum. *w_i_*= cumulative Akaike weight. “+” denotes additive effects and “×” denotes interaction effects. Models with ΔAICc ≤ 2 were considered to have substantial support. ^a^ Temporal trend. ^b^ Mean February temperatures on Central American nonbreeding grounds. ^c^ Mean January-February temperatures on Central American nonbreeding grounds. ^d^ Latitude bands (30°, 35°, 40°, and 45°N). ^e^ Mean June-October temperatures on North American breeding grounds.

| **Model** | ***k*** | **Dev.** | **$\Delta AICc$** | **$w_i$** | **Cum. $w_i$** |
| --- | --- | --- | --- | --- | --- |
| Year<sup>a</sup> | 3 | 9979.12 | 0.00 | 0.46 | 0.46 |
| Year + February temperature<sup>b</sup> | 4 | 9977.84 | 0.72 | 0.32 | 0.79 |
| Year + Winter temperature<sup>c</sup> | 4 | 9978.64 | 1.54 | 0.21 | 1.00 |
| Winter temperature | 3 | 10054.86 | 75.74 | 0.00 | 1.00 |
| February temperature | 3 | 10065.26 | 86.14 | 0.00 | 1.00 |

**Table 5.** Model selection results evaluating a priori hypotheses for predictors of migration
| **Model** | **k** | **Dev.** | **$\Delta\text{AICc}$** | **$w_i$** | **Cum. $w_i$** |
| --- | --- | --- | --- | --- | --- |
| Year × Latitude <sup>d</sup> | 5 | 44389.88 | 0.00 | 0.45 | 0.45 |
| Year + Latitude | 4 | 44392.44 | 0.55 | 0.34 | 0.79 |
| Fall temperature <sup>e</sup><br>+ Year ×<br>Latitude | 6 | 44389.42 | 1.54 | 0.21 | 1.00 |
| Year | 3 | 44767.40 | 373.50 | 0.00 | 1.00 |
| Fall temperature<br>+ Latitude | 4 | 45598.60 | 1206.70 | 0.00 | 1.00 |
| Fall temperature | 3 | 45952.46 | 1558.56 | 0.00 | 1.00 |
| Latitude | 3 | 46243.72 | 1849.83 | 0.00 | 1.00 |

Departure timing was best explained by models incorporating both year and latitude. The model including a year × latitude interaction received the strongest support, although an additive model containing both predictors was similarly well supported. These results indicate that departure phenology varies both temporally and across latitudes, while providing limited support for differences among latitudes in the rate or magnitude of temporal change. A third competitive model included fall temperature in addition to the interaction term. However, models including fall temperature alone performed poorly. Despite this, temperatures increased across both nonbreeding (Figure 4) and breeding (Figure 5) ranges over the study period.

**Figure 4.**
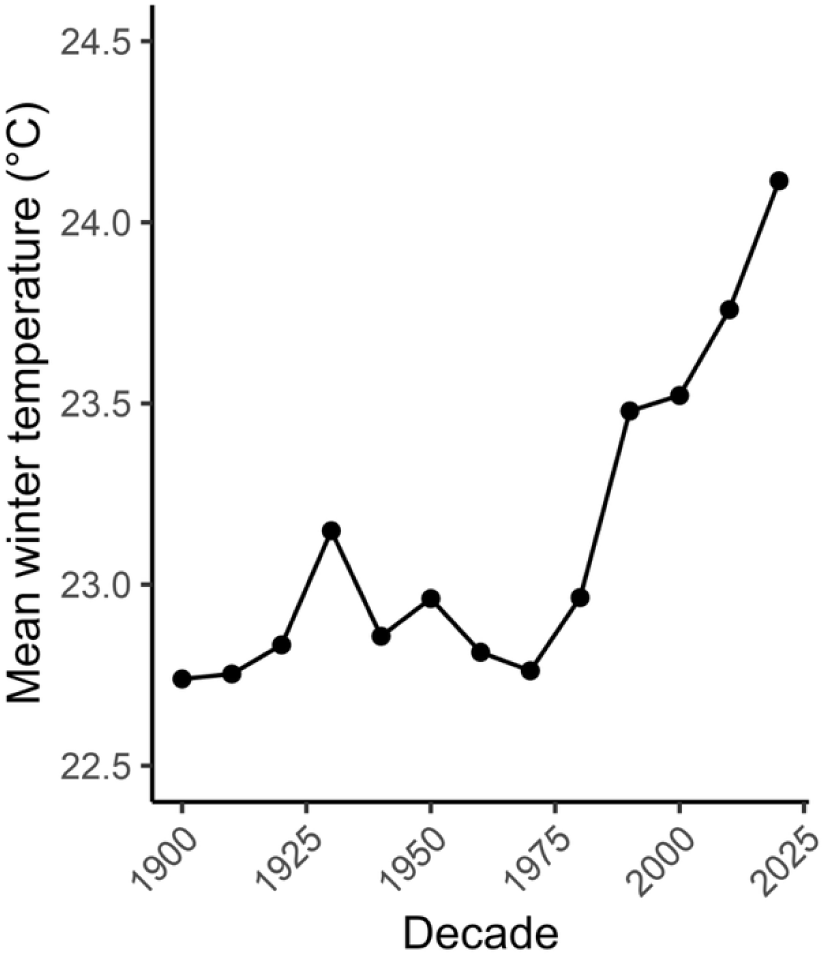
Average winter temperatures (January-February) in the nonbreeding range of *A. colubris* across decades. Alt text: Line graph depicting mean winter temperatures (°C) across decades 1900s to 2020s in the nonbreeding range of *A. colubris*. Temperatures are relatively stable through much of the 20th century, followed by a pronounced increase beginning in the 1980s and peaking in the most recent decades.

**Figure 5.**
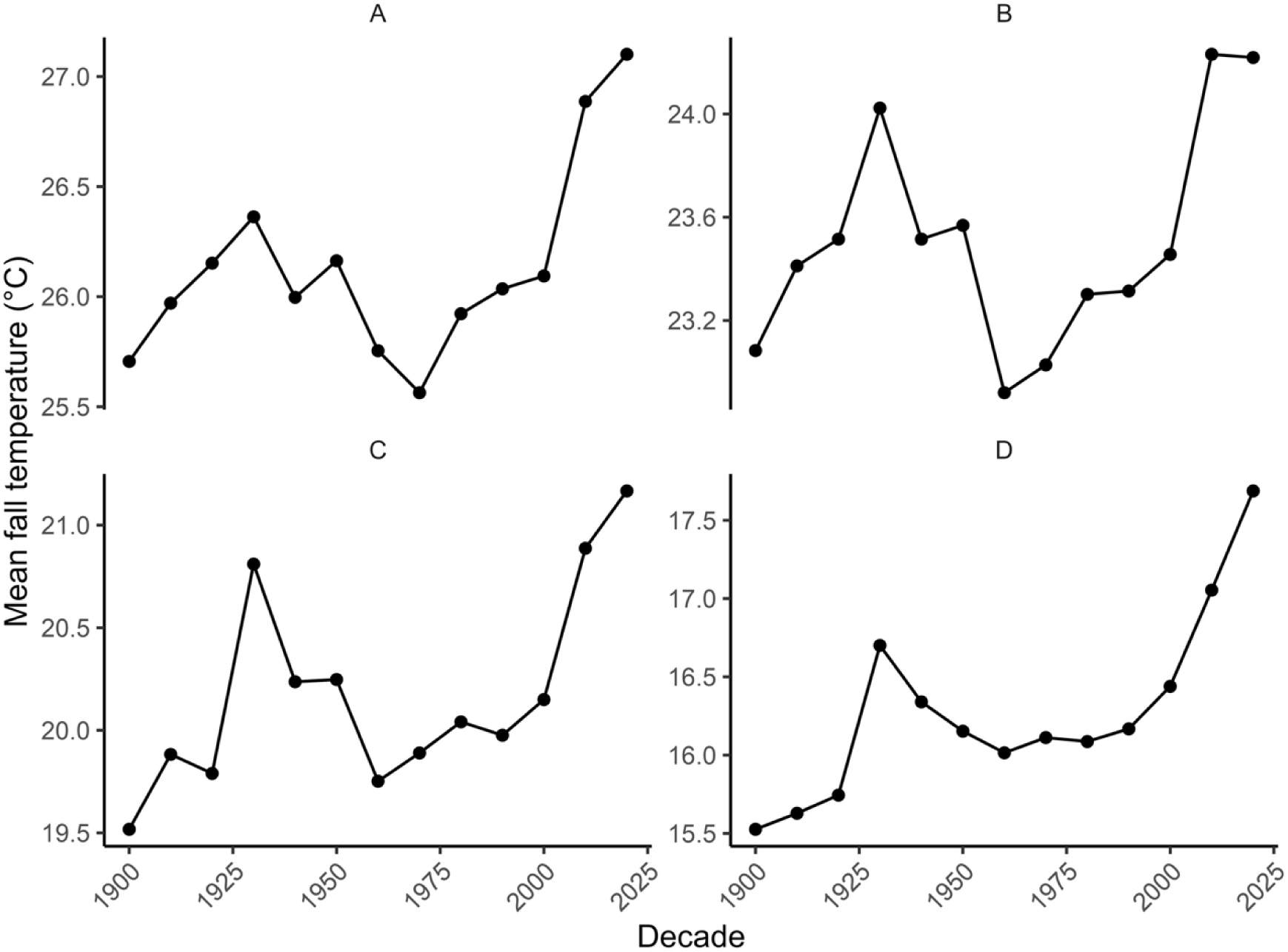
Average fall temperatures (June-October) in the breeding range of *A. colubris* by latitude bands across decades. (**A**) 30°N, (**B**) 35°N, (**C**) 40°N, and (**D**) 45°N. Alt text: 4-panel line graph showing mean fall temperature across decades 1900s to 2020s for 4 breeding latitude bands of *A. colubris.* There is noticeable warming after the 1980s, with the highest values occurring in recent decades.

## DISCUSSION

This study provides a broad-scale assessment of changes in *A. colubris* migration phenology in the United States over the past century. Mean first arrival dates at breeding areas have advanced significantly, with shifts ranging from 14.2 to 27.9 days earlier in recent years. These findings are consistent with previous studies on this species and reflect a global trend toward earlier spring migration in birds (Fernando and Gunawardena 2024). However, there have been no studies in the past decade reassessing changes in *A. colubris* migration. This may explain why past reports documented smaller changes in arrival dates, consistent with the observed decade-by-decade advancement in first arrival timing (Figure 1) (Gallinat et al. 2026), as well as the expansive spatial and temporal coverage of the present dataset that was not previously available. For example, Courter et al. (2013) reported a 14.3-day advancement at 35°N and a 13-day advancement at 40°N when comparing NABPP (1880-1969) and Journey North data (2001-2010), while the present study found 25- and 23.8-day shifts at the same latitudes. Similarly, Butler (2003) reported a 6.3-day shift in Worcester, Massachusetts (42°N, – 71°W) from 1932 to 1993, and Wilson et al. (2000) found a 4-day shift in Maine (44°N, 100°W) between the intervals 1899-1911 and 1994-1997. In contrast, this study identified 20.6- and 23.8- day shifts within the 40°N latitude band.

To our knowledge, no other study has assessed changes in fall migration timing for *A. colubris*. Although, given less studied than spring migration (Gallinat et al. 2015), numerous investigations have detected shifting departure dates from the United States in other birds. Reported trends vary by species, with suggested factors such as body measurements, migration distance, brood size, and diet proposed to influence the direction and magnitude of change (Jenni and Kéry 2003; Bitterlin and Van Buskirk 2014; Miles et al. 2017; Zimova et al. 2021). Furthermore, no single metric has been shown to fully capture patterns of changing phenology, and substantial uncertainty remains regarding interspecific differences in departure timing shifts (Bitterlin and Van Buskirk 2014; Miles et al. 2017). Nonetheless, the later departure dates observed in recent years in this study, ranging from 17.3 to 33.7 days, are consistent with delayed phenological shifts documented in other taxa (Bitterlin and Van Buskirk 2014; Zaifman et al. 2017).

Although climate change has been widely implicated in shifting migratory behavior in avian species (Usui et al. 2017; Zaifman et al. 2017; Cohen et al. 2018; Hinchcliffe and Tkaczynski 2025), the results of this study provide limited support for temperature as a primary driver of changes in *A. colubris* migration timing. However, the observed increases in temperature across both the nonbreeding and breeding ranges agree with the growing evidence of climatic warming during winter and fall seasons (Bevacqua et al. 2025). While photoperiod is regarded as the primary cue that triggers migration (Liddle et al. 2022), local temperature fine-tunes the timing of arrival and departure (Tottrup et al. 2010). It is therefore notable that increasing temperatures did not largely account for observed phenological shifts, particularly given that a previous study associated warmer February conditions in Central America with earlier spring arrival in this species (Courter et al. 2013). Differences between studies likely reflect both methodological and data-related factors. Courter et al. (2013) evaluated temperature effects using a single weather station and assessed statistical significance independently, whereas the present study utilized spatially extensive climate data and evaluated competing hypotheses using an information-theoretic model selection framework. In the present analysis, temperature variables alone received little support, and the inclusion of temperature alongside predictors resulted in only marginal improvements in model fit relative to simpler temporal models. Moreover, although winter temperatures increased across nonbreeding ranges over time, low variance inflation factors (VIF<2.5) indicated that multicollinearity was not severe in the arrival analyses. These findings suggest that temperature does not exert a strong independent influence on migration timing in *A. colubris*. Instead, broader temporal changes, such as other environmental cues or endogenous processes, are potentially contributing to the observed phenological shifts.

One potential mechanism may be the winter range of hummingbirds shifting northward into the southern United States due to the increase in predictable food sources through backyard bird feeding (Parmesan and Yohe 2003). A more northerly winter distribution would reduce the distance of spring migration, causing earlier arrival dates (Robb et al. 2008). In relation to the protraction of fall migration, delayed departure dates in other avian species have been attributed to re-nesting following failed breeding attempts (Zimova et al. 2021; Fernando and Gunawardena 2024). This behavior may reflect a response to mismatches between migration timing and peak food availability, which can reduce reproductive success, especially in species that also are arriving earlier in spring (Moller et al. 2010; Zimova et al. 2021). Postnuptial molt may also constrain departure timing, as molt is energetically demanding and must occur within the limited interval between breeding and migration (Tsuru and Tonra 2026). If re-nesting delays molt timing, molting could exacerbate delayed departure. Consistent with this possibility, fall migration timing has become more delayed in species that molt prior to migration, such as the *A. colubris* (Bitterlin and Van Buskirk 2014).

*A. colubris* spring migration pace advanced north of 40°N in both recent and historical periods (Table 2). Similar latitudinal effects on migration speed have been observed in other bird species and may reflect higher fuel deposition rates driven by increased primary productivity and longer daily foraging time at higher latitudes (Aharon-Rotman et al. 2016). Thus, birds may require shorter stopover durations. Minor differences observed in migration pace between contemporary data sources eBird and Journey North are likely a result of variation in methodology, with eBird offering broader spatial coverage and more frequent observations compared to the first arrival-based approach of Journey North. Limitations from the NABPP may also explain the discrepancy in fall migration pace between the 35-30°N latitude bands across data sources. During the historical period, it is likely that fewer observations were made at feeders, given that the use of supplemental sugar water has increased in recent decades (Robb et al. 2008), reducing the detection of birds. Modern arrivals are also reported online, which may increase observer vigilance when *A. colubris* are known to be present in the area (Backstrom et al. 2025). These considerations would influence both estimated migration pace and shifts in migration phenology (Koh and Opitz 2025). However, despite these limitations, we are confident that the results of this study provide meaningful biological insights into spatial and temporal changes in migration patterns, aligning with a growing body of evidence that birds are migrating earlier in spring and departing later in fall.

## Conclusion

Understanding species and ecosystem responses remains a central challenge in climate change research. This study demonstrates marked shifts in *A. colubris* migration phenology over time, with clear trends toward earlier spring arrivals and later fall departures to their breeding grounds in the eastern United States. However, increased temperatures were not significant independent predictors of these changes, possibly suggesting that other factors may be influencing migration timing. By improving our understanding of these dynamics, we can make informed conservation and policy decisions to protect migratory species under continued climate change.

