## Supplementary Materials Table S1 for "Advancement of spring arrival and delayed autumn departure in Ruby-throated Hummingbird (Archilochus colubris) migration phenology over more than a century"

Supplementary Tables to support results reported in preprint manuscript titled “Advancement of spring arrival and delayed autumn departure in Ruby-throated

Hummingbird (Archilochus colubris) migration phenology over more than a century” by Sydney M. Pierce and Terra M. Swartz

**Note:** All pairwise comparisons in Supplemental Tables S1-S11 were conducted using Tukey HSD tests following a two-way ANOVA. Positive values indicate later timing in the first-named source, while negative values represent later timing in the second-named source. *JN* = Journey North; *NABPP* = North American Bird Phenology Program.

**Table S1.** Differences in *A. colubris* mean first arrival dates among data sources, the historical NABPP and recent Journey North and eBird, averaged across all latitudes.

| **Comparison** | **Mean difference (days)** | **5% CI lower** | **95% CI upper** | **Adjusted *P*-value** |
| --- | --- | --- | --- | --- |
| JN vs. eBird | 0.0 | -0.5 | 0.5 | 0.99 |
| NABPP vs. eBird | 19.5 | 18.9 | 20.1 | <0.001 |
| NABPP vs. JN | 19.5 | 19.0 | 20.0 | <0.001 |

**Table S2.** Mean spring migration pace of *A. colubris* between successive 5° latitude bands for the recent eBird data source.

| **Latitude (°N)** | **Mean difference (days)** | **5% CI lower** | **95% CI upper** | **Adjusted *P*-value** |
| --- | --- | --- | --- | --- |
| 30 → 35 eBird | 21.4 | 19.2 | 23.7 | <0.001 |
| 35 → 40 eBird | 25.7 | 24.2 | 27.1 | <0.001 |
| 40 → 45 eBird | 11.7 | 10.3 | 13.1 | <0.001 |

**Table S3.** Mean spring migration pace of *A. colubris* between successive 5° latitude bands for the recent Journey North data source.

| **Latitude (°N)** | **Mean difference (days)** | **5% CI lower** | **95% CI upper** | **Adjusted *P*-value** |
| --- | --- | --- | --- | --- |
| 30 → 35 JN | 14.0 | 12.5 | 15.5 | <0.001 |
| 35 → 40 JN | 19.6 | 18.6 | 20.6 | <0.001 |
| 40 → 45 JN | 12.6 | 11.5 | 13.6 | <0.001 |

**Table S4.** Mean spring migration pace of *A. colubris* between successive 5° latitude bands for the historical NABPP data source.

| **Latitude (°N)** | **Mean difference (days)** | **5% CI lower** | **95% CI upper** | **Adjusted *P*-value** |
| --- | --- | --- | --- | --- |
| 30 → 35 NABPP | 22.1 | 18.1 | 26.0 | <0.001 |
| 35 → 40 NABPP | 18.4 | 16.3 | 20.4 | <0.001 |
| 40 → 45 NABPP | 5.4 | 4.1 | 6.7 | <0.001 |

**Table S5.** Differences in *A. colubris* mean first arrival day between recent data sources, Journey North and eBird, across four latitude bands.

| **Latitude (°N)** | **Mean difference (days)** | **5% CI lower** | **95% CI upper** | **Adjusted *P*-value** |
| --- | --- | --- | --- | --- |
| 30 JN vs. 30 eBird | 10.3 | 7.9 | 12.6 | <0.001 |
| 35 JN vs. 35 eBird | 2.8 | 1.5 | 4.2 | <0.001 |
| 40 JN vs. 40 eBird | -3.3 | -4.4 | -2.1 | <0.001 |
| 45 JN vs. 45 eBird | -2.4 | -3.7 | -1.0 | <0.001 |

**Table S6.** Differences in *A. colubris* mean first arrival day between data sources, the historical NABPP and recent eBird, across four latitude bands.

| **Latitude (°N)** | **Mean difference (days)** | **5% CI lower** | **95% CI upper** | **Adjusted *P*-value** |
| --- | --- | --- | --- | --- |
| 30 NABPP vs. 30 eBird | 27.2 | 23.3 | 31.2 | <0.001 |
| 35 NABPP vs. 35 eBird | 27.9 | 25.7 | 30.0 | <0.001 |
| 40 NABPP vs. 40 eBird | 20.6 | 19.3 | 21.8 | <0.001 |
| 45 NABPP vs. 45 eBird | 14.2 | 12.8 | 15.7 | <0.001 |

**Table S7.** Differences in *A. colubris* mean first arrival day between data sources, the historical NABPP and recent Journey North, across four latitude bands.

| **Latitude (°N)** | **Mean difference (days)** | **5% CI lower** | **95% CI upper** | **Adjusted *P*-value** |
| --- | --- | --- | --- | --- |
| 30 NABPP vs. 30 JN | 17.0 | 13.3 | 20.6 | <0.001 |
| 35 NABPP vs. 35 JN | 25.0 | 23.1 | 27.0 | <0.001 |
| 40 NABPP vs. 40 JN | 23.8 | 22.7 | 24.9 | <0.001 |
| 45 NABPP vs. 45 JN | 16.6 | 15.3 | 17.9 | <0.001 |

**Table S8.** Differences in *A. colubris* mean last departure dates between data sources, the historical NABPP and recent eBird, averaged across all latitudes.

| **Comparison** | **Mean difference (days)** | **5% CI lower** | **95% CI upper** | **Adjusted *P*-value** |
| --- | --- | --- | --- | --- |
| NABPP vs. eBird | -30.6 | -31.8 | -29.4 | <0.001 |

**Table S9.** Mean fall migration pace of *A. colubris* between successive 5° latitude bands for the recent eBird data source.

| **Latitude (°N)** | **Mean difference (days)** | **5% CI lower** | **95% CI upper** | **Adjusted *P*-value** |
| --- | --- | --- | --- | --- |
| 45 → 40 eBird | -8.4 | -11.1 | -5.7 | <0.001 |
| 40 → 35 eBird | -6.5 | -9.4 | -3.6 | <0.001 |
| 35 → 30 eBird | -2.6 | -8.3 | 3.0 | 0.84 |

**Table S10.** Mean fall migration pace of *A. colubris* between successive 5° latitude bands for the historical NABPP data source.

| **Latitude (°N)** | **Mean difference (days)** | **5% CI lower** | **95% CI upper** | **Adjusted *P*-value** |
| --- | --- | --- | --- | --- |
| 45 → 40 NABPP | -8.0 | -11.4 | -4.7 | <0.001 |
| 40 → 45 NABPP | -5.1 | -10.7 | 0.5 | 0.11 |
| 35 → 30 NABPP | -19.0 | -29.4 | -8.7 | <0.001 |

**Table S11.** Differences in mean last departure date of *A. colubris* between data sources, the historical NABPP and recent eBird, across four latitude bands.

| **Latitude (°N)** | **Mean difference (days)** | **5% CI lower** | **95% CI upper** | **Adjusted *P*-value** |
| --- | --- | --- | --- | --- |
| 30 NABPP vs. 30 eBird | -17.3 | -27.6 | -6.9 | <0.001 |
| 35 NABPP vs. 35 eBird | -33.7 | -39.3 | -28.0 | <0.001 |
| 40 NABPP vs. 40 eBird | -32.2 | -35.1 | -29.4 | <0.001 |
| 45 NABPP vs. 45 eBird | -31.8 | -35.0 | -28.7 | <0.001 |

**Note:** Model selection results for Supplemental Tables 12-13 were ranked using AICc. *k* = number of estimated parameters; ΔAICc = difference from the best-supported model; *w_i_* = Akaike weight; Cum. *w_i_* = cumulative Akaike weight; Dev. = deviance (−2lnL). “+” denotes additive effects and “×” denotes interaction effects. Models with ΔAICc ≤ 2 were considered to have substantial support.

**Table S12.** Model selection results for predictors of first arrival timing. Year represents temporal trends; February temperature refers to temperatures on Central American non‑breeding grounds; winter temperature represents mean January-February temperatures on the Central American non‑breeding grounds.

| **Model** | ***k*** | **AICc** | **ΔAICc** | ***w_i_*** | **Cum. *w_i_*** | **Log likelihood** | **Dev.** |
| --- | --- | --- | --- | --- | --- | --- | --- |
| Year | 3.00 | 9985.14 | 0.00 | 0.46 | 0.46 | -4989.56 | 9979.12 |
| Year + February temperature | 4.00 | 9985.86 | 0.72 | 0.32 | 0.79 | -4988.92 | 9977.84 |
| Year + Winter temperature | 4.00 | 9986.68 | 1.54 | 0.21 | 1.00 | -4989.32 | 9978.64 |
| Winter temperature | 3.00 | 10060.88 | 75.74 | 0.00 | 1.00 | -5027.43 | 10054.86 |
| February temperature | 3.00 | 10071.28 | 86.14 | 0.00 | 1.00 | -5032.63 | 10065.26 |
| Null | 2.00 | 10119.54 | 134.40 | 0.00 | 1.00 | -5057.76 | 10115.52 |

**Table S13.** Model selection results for predictors of last departure timing. Year represents temporal trends; fall temperature represents the mean June-October temperature on North American breeding grounds; latitude refers to latitudinal bands 30°, 35°, 40°, and 45°N.

| **Model** | ***k*** | **AICc** | **ΔAICc** | ***w_i_*** | **Cum. *w_i_*** | **Log likelihood** | **Dev.** |
| --- | --- | --- | --- | --- | --- | --- | --- |
| Year × Latitude | 5 | 44399.90 | 0.00 | 0.45 | 0.45 | -22194.94 | 44389.88 |
| Year + Latitude | 4 | 44400.45 | 0.55 | 0.34 | 0.79 | -22196.22 | 44392.44 |
| Fall temperature + Year × Latitude | 6 | 44401.44 | 1.54 | 0.21 | 1.00 | -22194.71 | 44389.42 |
| Year | 3 | 44773.41 | 373.50 | 0.00 | 1.00 | -22383.70 | 44767.40 |
| Fall temperature + Latitude | 4 | 45606.6 | 1206.70 | 0.00 | 1.00 | -22799.30 | 45598.60 |
| Fall temperature | 3 | 45958.46 | 1558.56 | 0.00 | 1.00 | -22976.23 | 45952.46 |
| Latitude | 3 | 46249.73 | 1849.83 | 0.00 | 1.00 | -23121.86 | 46243.72 |
| Null | 2 | 46715.09 | 2315.19 | 0.00 | 1.00 | -23355.55 | 46711.10 |

**Table S14.** Average winter temperatures (January-February) in the non-breeding range of *A. colubris* across decades.

| **Decade** | **Mean Temp. (°C)** |
| --- | --- |
| 1900 | 22.7 |
| 1910 | 22.8 |
| 1920 | 22.8 |
| 1930 | 23.1 |
| 1940 | 22.9 |
| 1950 | 23.0 |
| 1960 | 22.8 |
| 1970 | 22.8 |
| 1980 | 23.0 |
| 1990 | 23.5 |
| 2000 | 23.5 |
| 2010 | 23.8 |
| 2020 | 24.1 |

**Table S15.** Average fall temperatures (June-October) in the breeding range of *A. colubris* at the 30°N latitude band.

| **Decade** | **Mean Temp. (°C)** |
| --- | --- |
| 1900 | 25.7 |
| 1910 | 26.0 |
| 1920 | 26.2 |
| 1930 | 26.4 |
| 1940 | 26.0 |
| 1950 | 26.2 |
| 1960 | 25.8 |
| 1970 | 25.6 |
| 1980 | 25.9 |
| 1990 | 26.0 |
| 2000 | 26.1 |
| 2010 | 26.9 |
| 2020 | 27.1 |

**Table S16.** Average fall temperatures (June-October) in the breeding range of *A. colubris* at the 35°N latitude band.

| **Decade** | **Mean Temp. (°C)** |
| --- | --- |
| 1900 | 23.1 |
| 1910 | 23.4 |
| 1920 | 23.5 |
| 1930 | 24.0 |
| 1940 | 23.5 |
| 1950 | 23.6 |
| 1960 | 22.9 |
| 1970 | 23.0 |
| 1980 | 23.3 |
| 1990 | 23.3 |
| 2000 | 23.5 |
| 2010 | 24.2 |
| 2020 | 24.2 |

**Table S17.** Average fall temperatures (June-October) in the breeding range of *A. colubris* at the 40°N latitude band.

| **Decade** | **Mean Temp. (°C)** |
| --- | --- |
| 1900 | 19.5 |
| 1910 | 19.9 |
| 1920 | 19.8 |
| 1930 | 20.8 |
| 1940 | 20.2 |
| 1950 | 20.2 |
| 1960 | 19.8 |
| 1970 | 19.9 |
| 1980 | 20.0 |
| 1990 | 20.0 |
| 2000 | 20.2 |
| 2010 | 20.9 |
| 2020 | 21.2 |

**Table S18.** Average fall temperatures (June-October) in the breeding range of *A. colubris* at the 45°N latitude band.

| **Decade** | **Mean Temp. (°C)** |
| --- | --- |
| 1900 | 15.5 |
| 1910 | 15.6 |
| 1920 | 15.7 |
| 1930 | 16.7 |
| 1940 | 16.3 |
| 1950 | 16.2 |
| 1960 | 16.0 |
| 1970 | 16.1 |
| 1980 | 16.1 |
| 1990 | 16.2 |
| 2000 | 16.4 |
| 2010 | 17.1 |
| 2020 | 17.7 |
